# VHL ligand binding increases intracellular level of VHL

**DOI:** 10.1101/2021.04.12.439487

**Authors:** Julianty Frost, Sonia Rocha, Alessio Ciulli

## Abstract

The von–Hippel Lindau (VHL) protein is a tumour suppressor protein frequently mutated in the VHL disease, which functions as substrate recognition subunit of a Cul2 E3 ubiquitin ligase (CRL2^VHL^). CRL2^VHL^ plays an important role in oxygen sensing, by binding and targeting Hypoxia Inducible Factor-alpha subunits (HIF-alpha) for ubiquitination and degradation. VHL is also commonly hijacked by heterobifunctional degrader molecules known as proteolysis-targeting chimeras (PROTACs). In previous work we reported the structure-based design and functional characterisation of VHL inhibitors (VH032 and VH298) that induce the HIF response in cells. Here, we use unbiased quantitative mass spectrometry to identify the proteomic changes elicited by the VHL inhibitor and compare this to hypoxia or broad-spectrum prolyl-hydroxylase domain (PHD) enzyme inhibitor IOX2. Our results demonstrate the VHL inhibitor selectively activates the HIF response that is also present in the hypoxia- and IOX2-induced proteomic changes. Interestingly, VHL inhibitors were found to selectively upregulate a single protein, which is VHL itself. Our analysis revealed that this occurs via protein stabilisation of VHL isoforms and not via changes in transcript levels. Increased VHL levels upon VH298 treatment resulted in turn to reduced levels of HIF-1α protein. Our results demonstrate the high specificity of VHL inhibitors and suggest that use of these inhibitors would not produce overtly side effects due to prolonged HIF stabilisation. They also exemplify the concept that small-molecule binding induced protein stabilisation can increase protein levels inside cells.

## Introduction

The von Hippel–Lindau (VHL) tumour suppressor is a multi-subunit Cullin RING E3 ligase (CRL2^VHL^) – composed of Cullin2 as the central scaffold subunit, Rbx1 as RING subunit, ElonginB and ElonginC as adaptor subunits, and VHL as substrate recognition subunit (1,2). VHL functions to specifically bind hydroxylated HIF-1α, mediating polyubiquitination and subsequent targeting of HIF-1α for proteasomal degradation (3). Hypoxia-inducible factors (HIFs) are transcription factors that regulate the response to reduced oxygen availability termed hypoxia. HIF is composed of an oxygen-insensitive β-subunit (HIF-β) that is stable, and an oxygen-labile α-subunit, of which three isoforms are known: HIF-1α, HIF-2α and HIF-3α. The recognition and ubiquitination of HIF-α subunits by VHL is dependent on proline hydroxylation of the oxygen-dependent degradation domain of HIF-α, a post-translational modification mediated by 2-oxoglutarate, Iron (II) dioxygenases called Prolyl-hydroxylases (PHDs). Proline hydroxylation results in a high-affinity binding of HIF-α to VHL and thus subsequent ubiquitination and proteasomal degradation of HIF-α under normal oxygen conditions (3).

Mutations in human VHL result in a number of abnormalities, collectively called VHL disease when occurring in the germ line (4). However, VHL is also often loss in clear cell renal cell carcinoma (ccRCC) (5). The fact that VHL is a tumour suppressor protein, and that VHL loss is found in ccRCC tumours, have led to the hypothesis that VHL would not be a good target to inhibit and that chronic HIF stabilisation by VHL inhibitors might have detrimental side effects (6). Apart from HIF, other substrates for VHL have been postulated (7). These include Aurora A, ZHX2, NDRG3 and B-Myb (8-11). Understanding VHL functions and the pharmacology of HIF stabilisers and hypoxia inducers is thus important in a variety of physiological and pathological conditions (12).

We had previously developed several compounds, able to selectively bind to and inhibit VHL (13,14). Of these, VH298, is a potent chemical probe that triggers the hypoxic response via a different mechanism to other HIF stabilisers, i.e. by blocking the VHL:HIF-α protein–protein interaction downstream of HIF-α hydroxylation (13). We have also characterised the activity of a related compound VH032 (15), in its ability to induce the transcriptional response to hypoxia and how this compares to hypoxia (1% O_2_) or PHD inhibition using an unbiased RNA-sequencing approach (16). VH298 has also been used to trigger the hypoxic response in mice, demonstrating its appropriateness for *in vivo* applications (17,18). In distinct applications, VHL ligands are widely used as part of heterobifunctional degrader molecules known as proteolysis-targeting chimeras (PROTACs) (19-21). We and others have extensively demonstrated the use of VHL ligands VH032 and VH298 in PROTACs targeting BET proteins (22-24), protein kinases (25-27) amongst many other target proteins, including E3 ligases themselves as demonstrated by VHL homo-PROTACs (28).

Given the widespread use and applications of VHL ligands, inhibitors and VHL-based PROTACs, it is important to understand the effect of VHL inhibition to the intracellular proteome in an unbiased fashion. Here, we investigate how VHL inhibitors alter the proteome of cells, using quantitative TMT-labelling mass spectrometry. We compare these alterations to those occurring after hypoxia or exposure to a 2-oxoglutarate dioxygenase inhibitor, IOX2. We show that VH032 and its more potent related compound, VH298, increase VHL protein levels – an increase that is not observed following hypoxia or IOX2 treatment. Increases in VHL protein levels are due to stabilisation of specific VHL isoforms and not alteration of mRNA. VHL protein increases result in reduction of HIF levels following prolonged VH298 treatment in cells.

## Results

### Proteomic analysis of different HIF stabilising agents

We have recently published our unbiased mRNA-seq analysis of several HIF stabilising agents, including hypoxia, a broad-spectrum PHD inhibitor (IOX2) and a VHL inhibitor (VH032) (Figure 1A) (16). In addition, we had characterised the cellular responses to a more potent and specific VHL inhibitor called VH298 (13,14,16). To gain a better understanding of the effect of inhibition of VHL by the small molecule VHL inhibitor, global proteome analysis was performed in an unbiased manner. In a similar approach to our RNA-seq analysis, HeLa cells were treated with 250 µM VH032, comparing with vehicle control (1% DMSO), hypoxia (1% O_2_) and PHD inhibitor (250 µM IOX2). Enabled by the 10-plex tandem mass tags (TMTs)-labelling strategy, a total of four conditions were performed in duplicate with the treatment time of 24 h, a time point long enough to observe the global proteome changes induced by HIF and any other changes elicited by the VHL inhibitor (Figure 1B). By use of the mass spectrometry-based method, more than 8,043 proteins were identified with FDR < 0.01 and quantified (Table S1).

**Figure 1.**
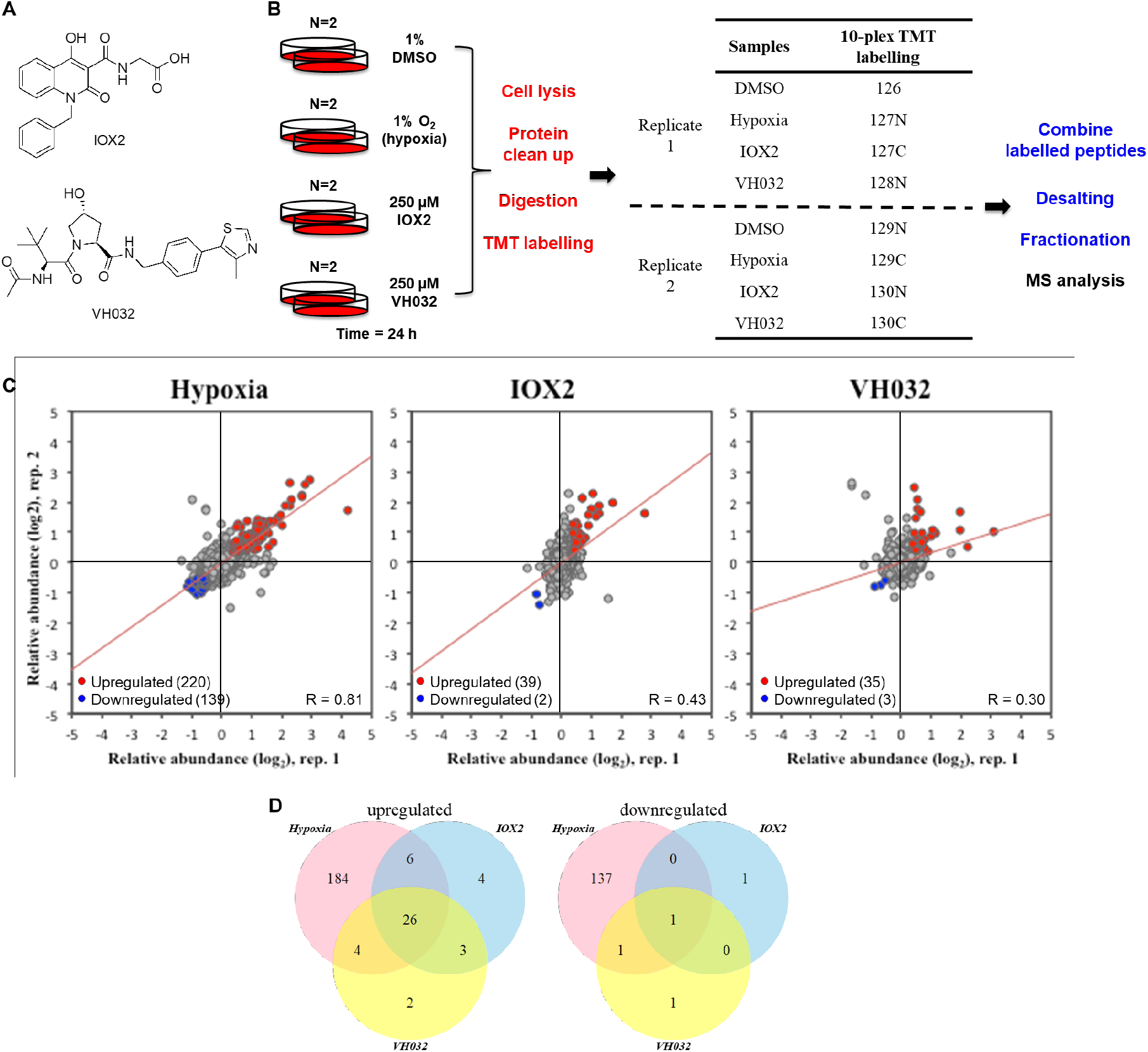
Unbiased proteomics identify VHL protein level increase upon VHL inhibitor treatment. (**A**) Chemical structures of IOX2 and VH032. (**B**) Diagram depicting the workflow of tandem mass tag (TMT)-labelling. HeLa cells were treated with 1% DMSO, 1% O2 (hypoxia), 250 μM IOX2, 250 μM VH032 for 24 h – in biological duplicate. Proteins were obtained by cell lysis, cleaned up, digested by trypsinisation, and labelled with 10-plex TMT labelling reagent. Labelled peptides were combined, desalted, fractionated and analysed by mass spectrometry (MS). Red text indicates steps carried out at the protein level, and blue are performed on peptides. (**C**) Scatter plot representation of relative protein abundances obtained for different treatment conditions compared to respective replicate of vehicle (DMSO)-treated cells, for a total of 8,043 proteins quantified. The two axes are relative abundance (log2FC) from two different replicates in this experiment. Red dots represent upregulated genes in both replicates (absolute fold change difference to DMSO > 1.3) and blue dots represent downregulated genes in both replicates (absolute fold change difference to DMSO < 0.7). Red line is the linear fit to the data. (**D**) Venn diagrams depicting the number of upregulated genes and downregulated genes in the two replicates comparing the presence of hypoxia, IOX2 or VH032 to DMSO control.

We next determined the reproducibility between the two replicates by plotting the relative abundance for each protein to respective DMSO control between the two replicates (Figure 1C). Hypoxia treatment showed the highest level of similarity in the proteins identified and quantified (Pearson correlation: 0.81), this was followed by IOX2 and VH032 treatments (Pearson correlation: 0.43 and 0.30, respectively). The mode of acquisition for the analysis is data dependent, which intrinsically generate variability within experiments (29). The lower correlation for the chemical inhibitors compared to hypoxia could be due to variability of inhibitor exposure across cells, but more work is needed to formally investigate this further. Analysis of the proteome changes identified revealed that the majority of upregulated proteins induced by VHL inhibitor were also identified in the hypoxia and IOX2 treatments, sharing 26 targets (Figure 1D, Table 1). Of these, 22 genes were also identified in our RNA-seq analysis, indicating changes to transcription vastly dependent on HIF (highlighted in Table 1). Of the additional 4 proteins shared by all of these stimuli, notably HIF-1α is present, and other proteins are CCKN (Cholecystokinin), RYR3 (Ryanodine Receptor 3), and CAR11 (Caspase Recruitment Domain Family Member 11).

**Table 1.**
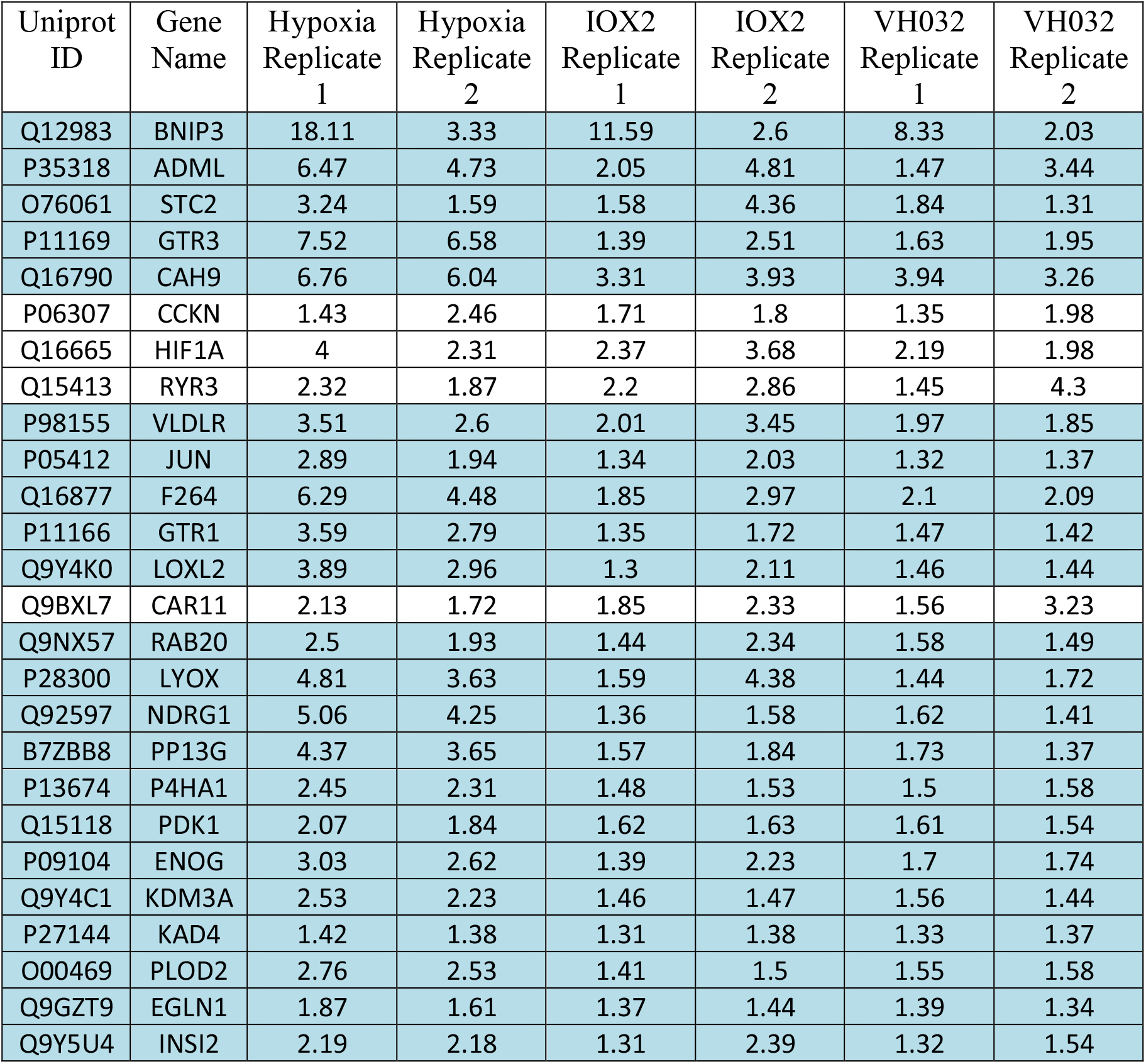
List of proteins upregulated in hypoxia, IOX2 and VH032 treatment. Proteins were selected at FDR < 0.01 and relative abundance to control DMSO > 1.3 for each replicate. Uniprot ID and gene name are listed with relative abundance to DMSO control. In blue highlight: transcripts of these proteins were found upregulated in hypoxia, IOX2 and VH032 by RNA-sequencing (16).

As expected, hypoxia induced the biggest changes in the proteome, including both induced and repressed protein expression (Figure 1D; Table S2). This was also reflected in the mRNA-seq analysis (16). In addition, more specific HIF inducers such as the VHL inhibitor and to some extent IOX2, broadly resulted in increased protein expression, with very few proteins found to be reduced in level (Figure 1D; Table S2). This is in line with the knowledge that HIF is associated with gene induction, and normally does not act as a transcriptional repressor (30).

### VHL inhibitors increase intracellular VHL protein levels

As we are particularly interested in the specificity of the VHL inhibitor, we turned our attention to the two proteins solely induced by VH032 compound: AMY1 (Amylase 1) and VHL. AMY1 is a protein previously shown to be induced by hypoxia in plants (31), so strictly not associated with VHL inhibition only. We therefore turned our attention to VHL itself. VHL protein abundance increased upon VHL inhibitor treatment, but not PHD inhibitor or hypoxia (Figure 2A-B). Relative to DMSO control, protein abundance of VHL increased to 1.59 for the first replicate and 1.53 for the second replicate (Figure 2A). In contrast, VHL protein abundance was unchanged in IOX2 (1.02 for replicate 1 and 0.94 for replicate 2) and decreased slightly in hypoxia (0.87 for replicate 1 and 0.89 for replicate 2).

**Figure 2.**
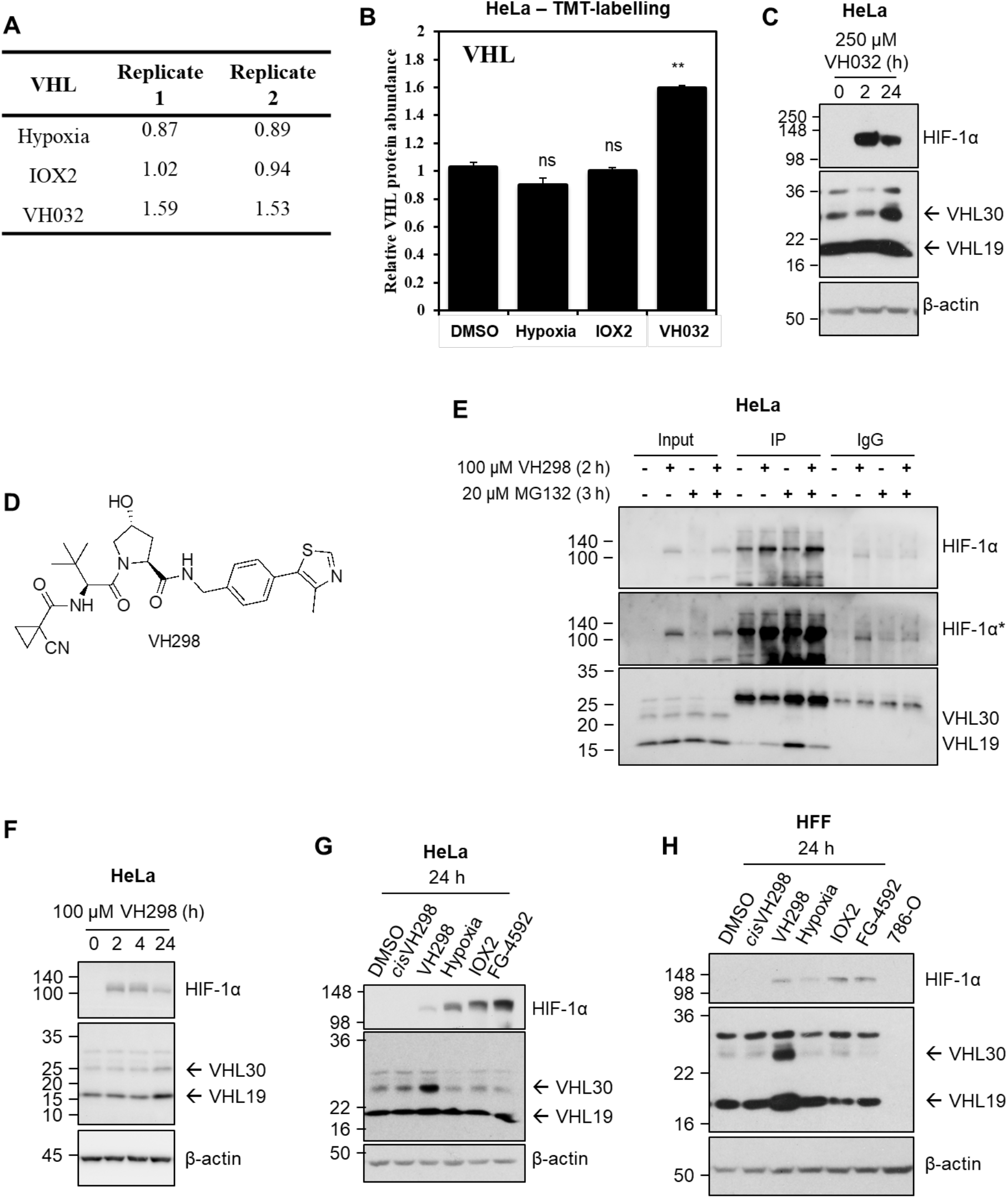
VHL protein levels increase in the presence of VHL inhibitors. (**A**) Relative abundance and (**B**) graph depicting relative VHL protein abundance with FDR < 0.01 comparing to DMSO control for hypoxia, IOX2 and VH032 treatments after 24 h. (**B**) Data are presented as means + SD from proteomic analysis (TMT-labelling) of two independent biological experiments. Two-tailed student’s t-test was performed to calculate p values, and levels of significance are denoted as follows: \**\*0*.*001<P*<0.01 and, ns: *P>0*.*05*. HeLa cells were treated with (**C**) 250 µM VH032 or (**F**) 100 µM VH298 for indicated time. (**D**) Chemical structure of VH298. (**E**) Co-immunoprecipitation on lysates from HeLa cells treated with vehicle DMSO (0.2% for 3 h), VH298 (100 μM for 2 h), MG132 (20 μM for 3 h), or VH298 and MG132 (100 μM VH298 for 2 h and 20 μM MG132 for 3 h) before lysis. 300 μg of protein were used to immunoprecipitate with the 2 μg HIF-1α antibody (Santa Cruz; sc-53546). Mouse immunoglobulin G (IgG; 2 μg) was used as a control. Inputs represent 10% of the starting material used per immunoprecipitation (IP). (**G**) HeLa or (**H**) HFF cells were treated with 1% DMSO, hypoxia (1% O2), and 100 μM of indicated compounds for 24 h. 786-O cell lysate was loaded in (**H**) as negative control for VHL bands. Protein levels were analysed by immunoblotting using antibodies against HIF-1α, VHL, and β-actin which acted as a loading control. HIF-1α* denotes longer exposure. The blots shown are representative of three independent experiments.

To validate the observation of the increase of VHL in the presence of VH032, we monitored VHL protein levels in HeLa cells treated with VH032 by western blot. VHL protein levels did not increase after a short treatment of 2 h, but showed marked increase in response to a longer treatment of 24 h with VH032 (Figure 2C). In the course of the investigation, we identified and characterised a more potent VHL inhibitor, with enhanced cell permeability and cellular activity (VH298; Figure 2D) that we qualified as a chemical probe (13,14). Inside cells, VH298 competed disrupted the binding between HIF-1α and VHL, as shown in co-immunoprecipitation experiments (Figure 2E). Therefore, we selected to use VH298 in our study moving forward. Like VH032, VH298 treatment resulted in increase in VHL protein levels (Figure 2F), also in a time-dependent manner, confirming that this increase is due to the inhibition of VHL by the small molecules. Additionally, the increase in VHL protein levels was not observed with the non-binding epimer *cis*VH298 (Figure 2G-H), further demonstrating that accumulation of VHL results specifically from small-molecule binding to VHL. VH298 also increased VHL protein levels in another cell context, HFF fibroblasts (Figure 2H). In both HeLa and HFF cells, the treatment of hypoxia or PHD inhibitor IOX2 did not increase VHL protein levels (Figure 2G-H), in agreement with the proteomic results (Figure 2A-B). We also confirmed that treatment with FG-4592, which is a more specific PHD inhibitor, showed similar results to those obtained with IOX2 – namely no alteration in VHL protein levels. Altogether, VHL protein levels were confirmed to be increased by the small molecule VHL inhibitors, not only in HeLa cells, in which the proteomic analysis was performed, but also in HFF fibroblasts, and this increase was not observed following hypoxia or in the presence of PHD inhibitors.

### Ligand-bound VHL increases protein stability

We next asked whether the increase in VHL protein levels in the presence of VHL inhibitor was due to the increase in mRNA levels. In qRT-PCR assays monitoring *VHL* mRNA, mRNA levels of *VHL* were not altered in the presence of VH032 or VH298 in HeLa (Figure 3A) or VH298 in HFF cells (Figure 3B). Similar to the unaltered VHL protein levels, hypoxia and IOX2 did not induce changes in *VHL* mRNA levels (Figure 3B). We next examined whether VHL might be auto-regulating itself to be degraded by the proteasome, similar to how VHL regulates the proteasomal degradation of HIF-1α. However, results showed that VHL protein levels did not increase in the presence of proteasome inhibitor MG132 (Figure 3C). We next asked whether the ligand-bound VHL might be more stable than the unbound form. We performed a cycloheximide chase experiment that inhibits *de novo* protein synthesis and monitored endogenous VHL protein levels to determine if the half-life of VHL increased in the presence of VHL inhibitor. Analysis revealed increased half-lives of both the long isoform of VHL, and the short isoform (Figure 3D-E). *VHL* encodes two major VHL isoforms: a 213 amino acids isoform (pVHL_1-213_), and a 160 amino acid isoform (pVHL_54-213_) (32,33). These two isoforms are also referred as pVHL30 and pVHL19, based on their apparent molecular masses upon protein electrophoresis. pVHL19 arises from an internal alternative translation initiation from Met54 within the open reading frame of VHL, and thereby missing the N-terminal pentameric acidic repeat domain (33,34). The VHL inhibitor VH298 increased the half-life of pVHL30 from 4 h to 6 h, while the half-life of VH298-bound pVHL19 increased to beyond the 6 h cycloheximide treatment. This result correlates to the increased protein levels for pVHL30 and pVHL19 seen in the presence of VHL inhibitors (Figure 2C,F-H). Taken together, this data indicates that the binding of VHL inhibitor leads to increased VHL protein levels, as a result of protein stabilisation, and delayed intracellular degradation.

**Figure 3.**
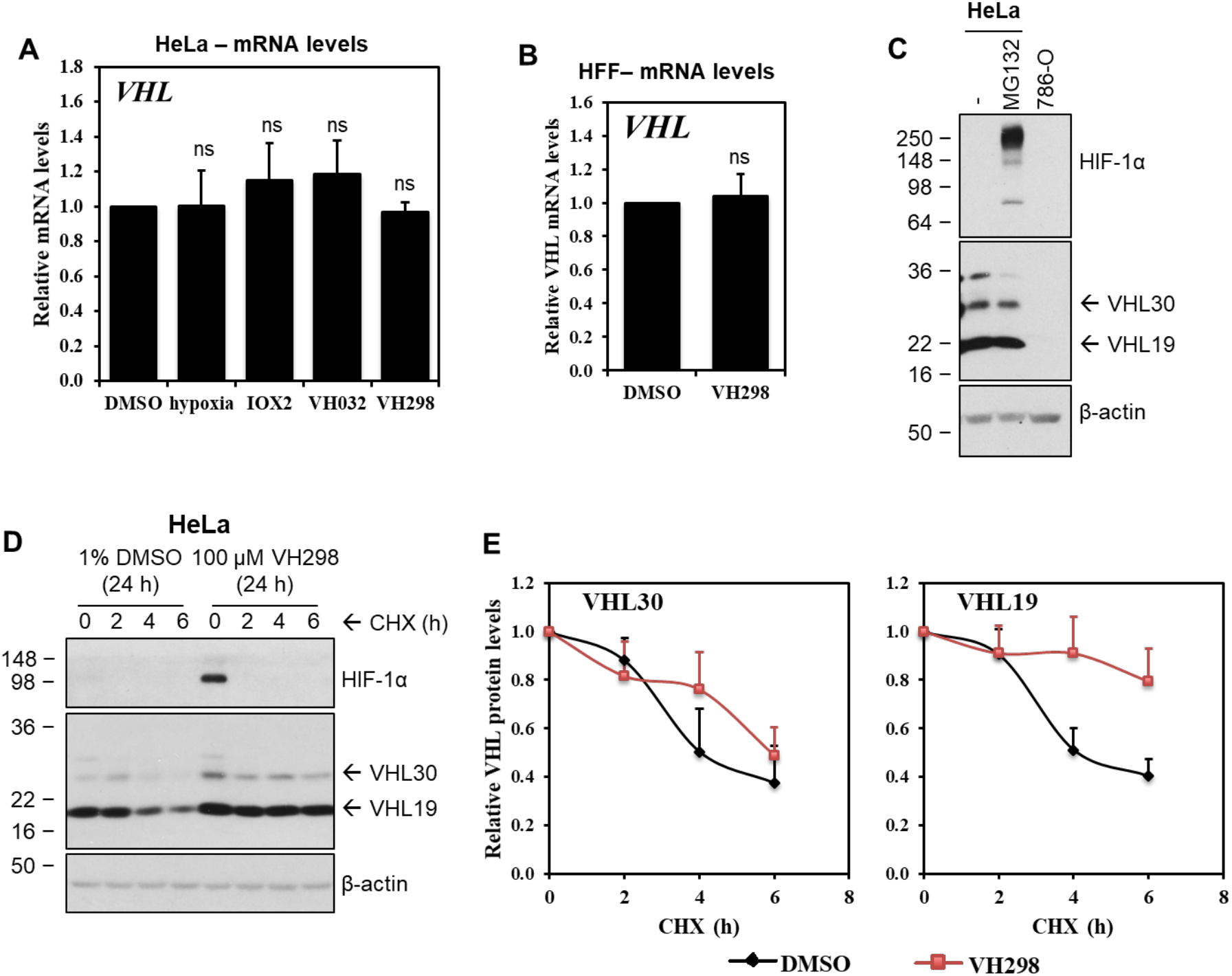
VHL inhibitor stabilizes VHL proteins at a post-translation level. VHL mRNA expressions in (**A**) HeLa cells treated with 1% DMSO, hypoxia (1% O2), and 250 μM IOX2, 250 µM VH032 or 100 µM VH298 for 16 h or in (**B**) HFF cells treated with 1% DMSO or 100 µM VH298 for 24 h. mRNA was collected, reverse transcribed and analysed by qRT-PCR. The shown levels of the indicated mRNAs were normalised to those of β-actin. Graphs depict the mean + SEM of three independent biological replicates. Two-tailed student’s t-test was performed to calculate p values, and levels of significance are denoted as follows: ns is *P>0*.*05*. (**C**) HeLa cells treated with 20 µM MG132 for 3 h. 786-O cell lysate was loaded as negative control for VHL and HIF-1α. (**D**) Half-life of VHL proteins of HeLa cells treated with DMSO negative control or VH298 VHL inhibitor was measured by treating cells with cycloheximide (CHX) for indicated times. (**E**) Protein levels of VHL30 and VH19 were quantified from multiple blots of different exposure time by ImageJ and plotted. Protein levels were analysed by immunoblotting using antibodies against HIF-1α, VHL, and β-actin, which acted as a loading control. The blots shown are representative of three independent experiments.

### The VH298-induced VHL enhances proteasomal degradation of HIF-1α in prolonged VH298 treatment

We have previously observed that HIF-1α protein levels are acutely upregulated in the presence of VH298, due to inhibition of the VHL–HIFα interaction, but this upregulation is not sustained in prolonged treatment, and eventually decreased over time (Figure 2F) (13). Since VHL protein levels increased in the presence of its inhibitor, we hypothesise that the decrease of HIF-1α levels observed in prolonged treatment of VHL inhibitor could be due to the increase of VHL protein levels. First, we investigated whether we could rescue the degraded HIF-1α by adding more VH298 to further inhibit the stabilised VHL. We determined the optimum concentration of VH298 to induce the maximum level of HIF-1α and the doses were found to be 400 µM and 100 µM in HeLa and HFF cells, respectively (Figure 4A). The optimum concentration of VH298 was then added to respective cells that had been treated with 100 µM VH298 for 24 h, at which point HIF-1α protein levels would have decreased. HIF-1α protein levels were monitored 2 h after the subsequent addition of VH298 (Figure 4B). Although HIF-1α increased slightly, the addition of more VH298 did not rescue the degraded HIF-1α to the same level as the initially stabilised HIF-1α at 2 h in both HeLa and HFF (Figure 4B). These results could indicate that the decrease of HIF-1α levels in prolonged VH298 treatment was not due to insufficient VH298 to inhibit VHL. However, it is also possible that the extra-added inhibitor may be limited by cell permeability and so may still be insufficient to fully saturate intracellular VHL, particularly given the significantly increased VHL levels under those conditions. Therefore, we next investigated whether the VH298-induced HIF-1α is degraded in a VHL-dependent manner by performing siRNA-mediated knockdown of VHL in HeLa cells, followed by VH298 treatment. In control siRNA-treated cells, VH298 treatment gradually resulted in reduction of HIF-1α levels, as expected. In contrast, under VHL knockdown by siRNA, HIF-1α remained stabilised over the entire course of 24 h compound treatment (Figure 4C). This suggests that the degradation of VH298-induced HIF-1α was indeed mediated by VHL. Interestingly, siRNA knockdown of VHL alone led to almost undetectable stabilisation of HIF-1α (Figure 4C-D), as previously shown (28). This is consistent with siRNA knockdown not being able to completely remove all endogenous VHL, and confirms that even very low amount of the remaining VHL appears to be sufficient to efficiently polyubiquitinate HIF-1α for proteasomal degradation.

**Figure 4.**
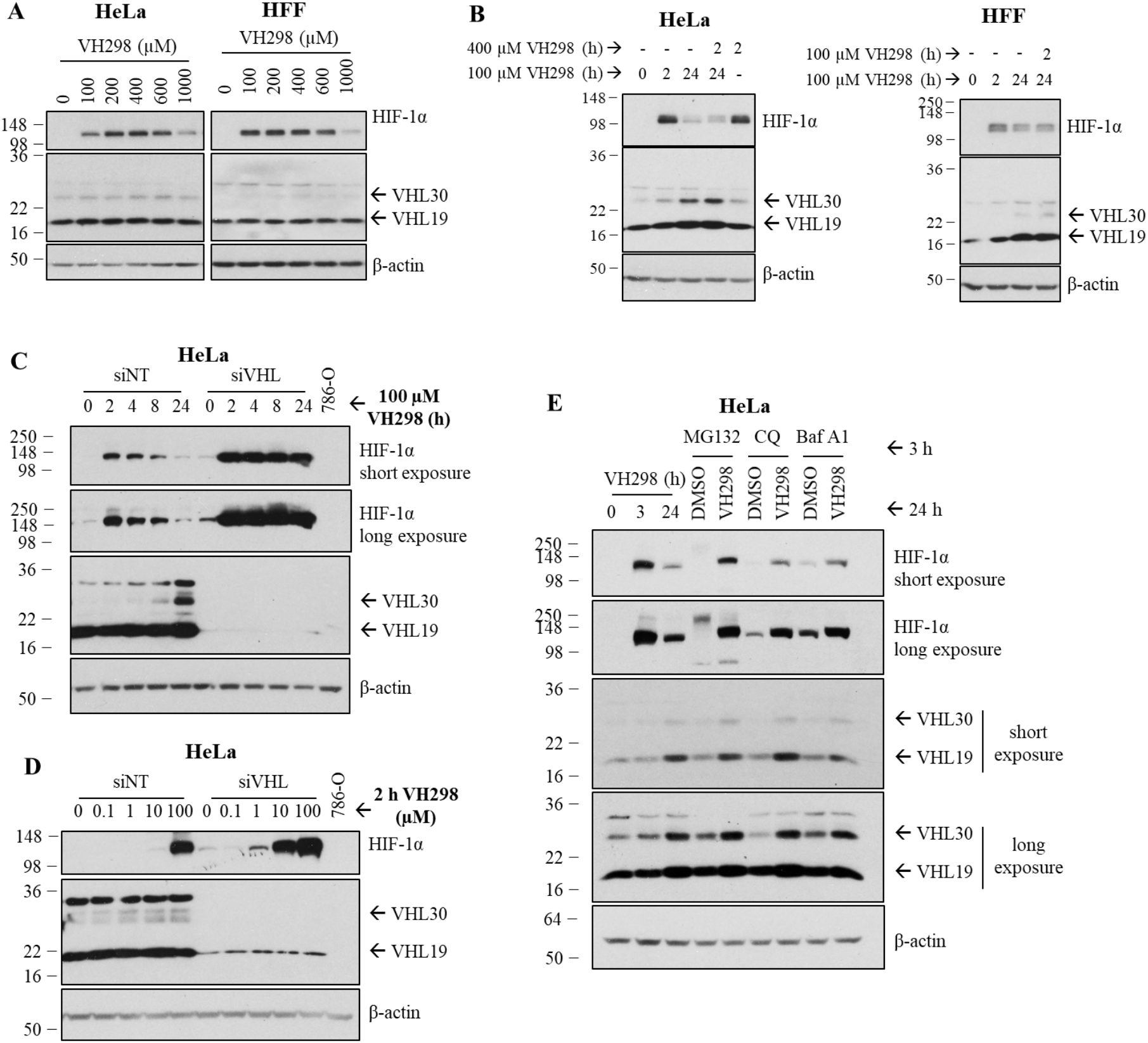
The decrease of HIF-1α protein levels in prolonged VH298 treatment is mediated by proteasomal degradation in a VHL-dependent manner. (A) Dose-dependent immunoblots of HIF 1α in HeLa (left panel) and HFF (right panel) cells treated with VH298 for 2 h. (**B**)HeLa and HFF cells were treated with 100 µM VH298 for 2 and 24 h. After 24 h treatment of 100 µM VH298, 400 µM VH298 was introduced to HeLa cells for 2 h and 100 µM VH298 to HFF cells for 2 h. A 2 h treatment with 400 µM VH298 was also included in HeLa cells. (**C-D**) HeLa cells were transfected with non-targeting siRNA control (siNT) or VHL siRNA (siVHL). (**C**) After 24 h, media was changed and transfected cells were treated with 100 µM VH298 for indicated times. (**D**) After 46 h, media was changed and transfected cells were treated with indicated concentrations of VH298 for 2 h. Cell lysate of 786-O was included as negative control for VHL. (**E**) HeLa cells were treated with 3 and 24 h of 100 µM VH298. After 24 h treatments of 1% DMSO or 100 µM VH298, cells were treated with proteasome inhibitor MG132 (20 µM), autophagy inhibitors chloroquine (CQ; 50 µM) or bafilomycin A1 (Baf A1; 50 nM) for 3 h. Protein levels were analysed by immunoblotting using antibodies against HIF-1α, VHL, and β-actin, which acted as a loading control. The blots shown are representative of three independent experiments.

We postulate that the high activity and catalytic efficiency of VHL might explain the requirement of relatively high concentration of VHL inhibitor in order to observe detectable levels of HIF-1α. To examine this hypothesis, a dose-dependent VH298 treatment was performed with and without siRNA-mediated VHL knockdown (Figure 4D). In the background of siRNA-mediated VHL knockdown, HIF-1α levels could be detected already at 1 µM concentration of VH298, in stark contrast to the negative control (siNT) background where HIF-1α levels were undetected at the same concentration of VH298, nor at 10-times higher concentration of VH298 (Figure 4D). Remarkably, a 10 µM inhibitor treatment under VHL knockdown was found to achieve similar HIF-1α stabilisation as 100 µM VH298 in the negative control siRNA background (Figure 4D). The combination of VH298 and the knockdown of VHL showed additive effect on accumulating HIF-1α (Figure 4D). Together, these data indicate that following treatment with VHL inhibitor VH298, HIF-1α is initially stabilised, and then eventually degraded in a VHL-dependent manner, as VHL level increase.

HIF-1α is known to be degraded via the proteasomal (35), as well as lysosomal (36) pathways. To investigate the pathway involved in the degradation of VH298-induced HIF-1α, HeLa cells were first treated with VH298 for 24 h, followed by proteasome inhibitor (MG132) or lysosome inhibitors (chloroquine or bafilomycin A1) for the last 3 h. As expected, HIF-1α was stabilised upon each of the three inhibitors (Figure 4E). After 24 h of VH298 treatment, MG132 rescued the degraded HIF-1α to similar protein levels as 3 h treatment of VH298 (Figure 4E). In contrast, lysosomal inhibitors were not as efficient; both chloroquine and bafilomycin A1 increased HIF-1α to a much lower extent compared to MG132. Altogether, these data indicate that in prolonged VH298 treatment, the HIF-1α proteins that accumulated in the presence of VHL inhibitor VH298 are eventually degraded mainly via the proteasomal pathway, in a VHL-dependent manner.

## Discussion

In this study we investigated the selectivity of a VHL inhibitor VH032 at the proteomic level. Using quantitative TMT-labelling based mass spectrometry analysis, VH032 effects were compared to those elicited by hypoxia and the broad-spectrum inhibitor IOX2. This analysis complemented our previous mRNA-seq analysis (16), as proteins that we identified changing were also captured at the mRNA level. This is important, as several global omics studies have reported a poor correlation of mRNA to protein (37). Furthermore, some transcripts are induced at different stages of the hypoxia response, and protein translation and stability might also be altered in hypoxia. Finally, we wanted to determine if the VHL inhibitor would have off-target effects at the protein level, not identified at the mRNA level. Our analysis confirmed that all HIF stabilising agents used share an overlap of proteins that are upregulated. The vast majority of proteins identified as induced in our analysis overlapped with our mRNA-seq as predicted. Furthermore, from the additional proteins identified, HIF-1α and known HIF targets were also present. This suggests that some HIF targets are induced at different times in the hypoxia response and that a more dynamic analysis of mRNA and protein would be required to fully cover all proteins induced by the stimuli used. Although VHL has been shown to control NF-KB activity (38-40), VH032 was found to target only the HIF response, as no NF-KB dependent signature was identified in our proteomic dataset or indeed in our RNA-seq dataset (16). This further supports the specificity of the VHL inhibitor used in this study.

Given the widespread use of VHL inhibitors and VHL-based PROTACs as chemical tools to study biology, and as therapeutics, we were particularly interested in assessing target-specific effects, and potentially identify any off-target effects. Our proteomics analysis revealed that VH032 is indeed exquisitely specific and selective for VHL. In fact, the only additional protein upregulated by VH032, not previously shown to be induced by hypoxia was VHL itself, as AMY1 had previously been identified as a hypoxia inducible protein (31). Stabilisation of VHL was also confirmed by using the more potent inhibitor VH298. Further analysis revealed that this is dependent on VHL isoform stabilisation and not alterations in mRNA levels. We have previously demonstrated direct binding of VH298 to VHL, which *in vitro* resulted in increased thermal stability both biophysically on the recombinant protein (41) as well as in a cellular environment (13). Interestingly, our results suggest that increased levels of VHL are responsible for reduced HIF-1α levels observed following prolonged exposure to VH298. Our data suggests that lowering the levels of VHL is able to circumvent the reduction of HIF-1α.

There are several implications for these observations. First, from a biological standpoint it suggests that the HIF system explores increased VHL protein level as yet another negative feedback loop in place to prevent excess levels of the HIF transcription factor for prolonged periods of time. This adds to the known feedback loop of increased PHD2 and PHD3 protein levels (42). Second, it implies that changes to VHL levels can in itself regulate HIF-1α levels in cells. This is a concept that needs further exploring. Although proline hydroxylation is in most cases sufficient to maintain intracellular HIF-1α at low level (3), increasing VHL levels can provide an additional mechanism to further reduce HIF-1α levels where necessary. Thus far, this mechanism remains understudied, and thus future research is needed to investigate this further.

Our observations are also relevant to the pharmacology of VHL inhibitors such as VH298 or its derivatives, as well as VHL-based PROTAC degraders. This is important given the wide use of these compounds as tools to study biology and the interest in their therapeutic development. VH298 has been shown to improve wound healing (18) and also tendon healing (17) in a rat model of injury. Other potential uses could be anaemia and mitochondrial dysfunction, where HIF stabilisation has been shown to be of benefit (43,44). Our results demonstrating high target specificity of VH298 and feedback leading to reduced HIF-1α levels upon prolonged inhibitor treatment suggest that the use of a medicinal derivative of this chemical probe in these conditions could be highly recommended, as it would be predicted to have low side effects in its response. However, further analysis is now needed *in vivo* in these pathological conditions to firmly establish this possibility. With regards to PROTACs, to our knowledge no study to date have reported of increased VHL levels upon PROTAC treatment. This may be in part due to the fact that VHL protein levels tend not to be monitored during PROTAC treatment, with the exception of homo-PROTACs and hetero-PROTACs aimed at inducing E3 ligase degradation (28,45). It is also conceivable to imagine that VHL stabilisation will not usually be observed due to the sub-stoichiometric mode of action of PROTACs, working at sufficiently low concentration to induce target protein degradation, as opposed to the occupancy-based mode of action of inhibitors (19). Nonetheless, we advocate for the field to monitor intracellular protein levels of VHL when using VHL-based PROTACs. Interestingly, low intracellular expression of the E3 ligase VHL in platelets has been invoked to explain and exploit the observed reduced dose-limiting toxicity of VHL-based Bcl-xL PROTAC degraders (46).

We exemplify a case where a potent protein-protein interaction inhibitor can increase the intracellular levels of its target protein, while leading to a decrease in substrate recognition and hence enzyme activity. While this may not be unprecedented, we were able to find only one other such example in the literature: the enzyme thymidylate synthase, which is induced upon stabilization by its inhibitor 5-fluoro-2′-deoxyuridine 5′-monophosphate (47). When inhibiting metabolic or signalling pathways, enzyme target inhibition is often coupled to a feedback change in metabolite or substrate/product level, which in turn increases the transcription and/or the translation of the gene encoding for the enzyme target (48,49). The net result is an increase in total target protein, leading to inhibitor/drug resistance. Other mechanisms of increase target level occur as a result of gene amplification (50), or release of the enzyme from binding to its own mRNA (51). Here, our results exclude transcriptional/translational upregulation, and instead indicate that protein stabilisation is the primary mechanism responsible for the increased intracellular levels of VHL. The finding that VHL inhibitors stabilize their target protein is consistent with current understanding of VHL structure and function. It is well documented that VHL acts as a tumour suppressor (6), and that the VHL protein has both moderate thermal stability (T_m_ ∼ 47 °C, see refs. (13) and (41)) and moderate cellular half-life (∼4-6 h, see Figure 3E herein). We therefore hypothesize that VHL stabilization upon small-molecule binding leads to a reduction in intracellular degradation of VHL. We speculate that such protection away from degradation could be the result of oligomerisation (to itself or partner proteins), by sequestration to other cellular compartments, or by stabilizing flexible, disordered regions on the VHL protein that normally confer its instability. Elucidating the detailed mechanism by which VHL inhibitors stabilise VHL warrants further mechanistic investigation in the future.

In conclusion, this work demonstrates the high specificity of VHL inhibitors VH032 and VH298 and that these inhibitors increase intracellular levels of their own target protein VHL, as a result of protein binding and stabilisation. This has important implications for the pharmacology of VHL inhibitors and potentially VHL-based PROTAC degraders. We suggest that chronic long-term use of VHL inhibitors might benefit from our newly identified VHL stabilisation, and therefore pharmacological VHL blockade might not be as detrimental to cells or organisms as previously thought.

## Experimental procedures

### Cell culture and hypoxia induction

Human cervix carcinoma cells HeLa and human foreskin fibroblasts HFF were obtained from American Type Culture Collection (ATCC), and propagated in DMEM supplemented with 10% fetal bovine serum (FBS), L-glutamine, 100 μg/mL of penicillin/streptomycin) at 37 °C. Cell lines were routinely tested for mycoplasma contamination using MycoAlert™ kit from Lonza.

### Hypoxia treatment

For hypoxia induction, cells were incubated at 1% O_2_ in an InVIVO 300 hypoxia workstation (Ruskin Technologies). In order to prevent reoxygenation, cells were lysed for protein extraction in the hypoxia workstation.

### Compound treatments

DMSO was used as vehicle control. The proteasome inhibitor MG132 was obtained from Merck/Millipore and used at the final concentration of 20 µM for 3 h. PHD inhibitors IOX2 and FG-4592 were purchased from Selleckchem. VHL inhibitors VH032, VH298 and non-binding epimer *cis*VH298 were synthesised by a group member of our laboratory (13,14). Compounds were added to cells for indicated length of time. Chloroquine (Merck) and baflomycin A1 (Selleckchem) were added to cells for 3 h at 50 µM and 50 nM, respectively.

### siRNA transfections

Small interfering RNA oligonucleotides were purchased from Eurofins/MWG, and used in a stock concentration of 20 μM. siRNAs were transfected using Interferin from Polyplus according to the manufacturer’s instructions. The oligonucleotide sequences used for siRNA knockdown are the following:

**siNT** (non-targeting): 5’ – AAC AGU CGC GUU UGC GAC UGG – 3’

**siVHL:** 5’ – GGA GCG CAU UGC ACA UCA ACG– 3’

### Immunoblotting

Cells were lysed in RIPA buffer (50 mM Tris pH 8, 150 mM NaCl, 1% NP-40, 0.5% sodium deoxycholate, 0.1% SDS, 250 M Na_3_VO_4_, 10 mM NaF, and a protease inhibitor cocktail (Roche) per 10 mL buffer. Proteins were resolved using sodium dodecyl sulfate polyacrylamide gel electrophoresis (SDS–PAGE), transferred onto polyvinylidene difluoride membranes in a semi-dry transfer system (BIORAD) and detected using primary antibodies, with β-actin as loading controls in mammalian cells.

Primary antibodies were used at following dilutions for mammalian cells: anti-HIF-1α (BD Biosciences; 610958; 1:1,000), anti-VHL (Cell Signalling; #2738; 1:1,000; product is discontinued), anti-VHL (Cell Signaling Technology; #68547; 1:1,000) and anti-β-actin (Cell Signaling Technology; #3700s; 1:10,000). Following incubation with a horseradish peroxidase-conjugated secondary antibody (Cell Signaling), chemiluminescence (Pierce) was used for immunodetection.

### Co-Immunoprecipitation

HeLa cells were lysed in RIPA buffer as above. 300 μg of cell lysates were used per immunoprecipitation condition. Protein lysates were incubated with 2 μg of HIF-1α antibody (Santa Cruz; sc-53546) or 2 μg of mouse IgG control antibody (Sigma) in a rotating platform at 4 °C overnight. 20 μl of packed protein-G-sepharose beads (Pierce) were used to recover the immuno-complexes, by incubation in a rotating platform for 1.5 h at 4 °C. Beads were washed with 1 × PBS buffer thrice. The complexes were eluted from beads with SDS loading buffer and resolved as described above by immunoblotting.

### Quantitative real-time PCR

Mammalian cells: RNA was extracted using the RNeasy Mini Kit (Qiagen) according to manufacturer’s protocol. RNA was reverse transcribed using the iScript™ cDNA Synthesis Kit (BIO-RAD). Real-time PCR was performed in triplicates using PerfeCTa® SYBR® Green FastMix® (Quanta Biosciences) in C1000 Touch Thermal Cycler (Bio-Rad Laboratories). *m*RNA levels were calculated based on averaged CT values and normalised to β-actin mRNA levels.

Primer sequences used are listed.

**VHL** forward, 5’ – CCT TGG CTC TTC AGA GAT G – 3’
**VHL** reverse, 5’ – TGA CGA TGT CCA GTC TCC T – 3’
**β-actin** forward, 5’ – CTC TTC CAG CCT TCC TTC CTG – 3’
**β-actin** reverse, 5’– GAA GCA TTT GCG GTG GAC GAT – 3’

### Protein sample preparation and tandem mass tag (TMT) labelling mass spectrometry

HeLa cells were treated with 0.05% DMSO, hypoxia (1% O_2_), 250 µM IOX2 and 250 µM VH032 for 24 hours. HeLa cells were lysed in 4% SDS in 100 mM Tris-HCl, pH 8.5 and 1x protease inhibitor (one cocktail tablet [Roche] per 10 mL of lysis buffer). Lysates were first sonicated for 6 cycles of 30 sec on/30 sec off under low power setting with Bioruptor™ Twin (Diagenode), and then centrifuged at 17,000 × g for 10 min at 4 °C. Samples could be stored at ?80 °C freezer. The experiments were performed in two biological replicates. The protein samples were further processed, including reduction, alkylation, digestion, desalting, TMT-labelling and fractionation according to previously described (24).TMT 10plex™ Isobaric Label Reagent Set (Thermo Fisher Scientific) was used despite two biological replicates of four conditions (a total of eight samples), as a duplicate of another condition was included in the mass spectrometry that was not part of this project. The forth duplicate treatment was VHL inhibitor VH101 which however led to confounding the results due to its cytotoxicity, so is not included (14). Samples were labelled according to indicated in Figure 1A.

### *nLC MS/MS* analysis

Analysis of peptides was performed on a Q Exactive™ HF Hybrid Quadrupole-Orbitrap™ Mass Spectrometer (Thermo Scientific) coupled with a Dionex UltiMate 3000 RSLCnano Systems (Thermo Scientific). The peptides from each fraction were separated using a mix of buffer A (2% acetonitrile and 0.1% formic acid in Milli-Q water (v/v)) and buffer B (80% acetonitrile and 0.08% formic acid in Milli-Q water (v/v)). The peptides were eluted from the column at a constant flow rate of 300 nl/min with a linear gradient from 95% buffer A to 40% buffer B in 122 min, and then to 98% buffer B by 132 min. The column was then washed with 95% buffer B for 15 min and re-equilibrated in 98% buffer A for 32 min. The Q Exactive™ HF Hybrid Quadrupole-Orbitrap™ Mass Spectrometer was used in data dependent mode. A scan cycle comprised MS1 scan (m/z range from 335-1800, with a maximum ion injection time of 50 msec, a resolution of 120 000 and automatic gain control (AGC) value of 3×10^6^) followed by 15 sequential dependant MS2 scans (with an isolation window set to 1.2 Da, resolution at 60000, maximum ion injection time at 200 msec and AGC 1×10^5^. To ensure mass accuracy, the mass spectrometer was calibrated on the first day that the runs are performed.

### Data processing and database searching

Raw MS data from 22 fractions were searched against Swiss-Prot database (8^th^ March 2015; *Homo sapiens*; 54,7964 sequences – after human taxonomy filter applied – 20,203) using the Mascot search engine (Matrix Science, Version 2.2) through proteome discovery software (version 1.4). Mascot significance threshold was set to 0.05. Trypsin/P was specified as the cleavage enzyme allowing up to two missed cleavages. Parameters for database search were as follow MS1 Tolerance: 10 ppm, MS2 Tolerance: 0.06 Da, fixed modification: Carbamidomethyl (C). Variable Modification: Oxidation (M), Dioxidation (M), Acetyl (N-term), Gln->pyro-Glu (N-term Q), hydroxyl (P), TMT10plex (N-term) and TMT10plex (K). The data was filtered by applying a 1% false discovery rate. Quantified proteins were filtered if the absolute fold change difference between conditions to respective DMSO of the same biological replicate was ≥ 1.3 for upregulated proteins and ≤ 0.7 for downregulated proteins.

## Data availability

Data supporting the findings of this study are available within the article (and its Supplementary Information files) and from the corresponding author upon reasonable request. The mass spectrometry proteomics data have been deposited to the ProteomeXchange Consortium via the PRIDE partner repository with the dataset identifier PXD025743.

## Supporting information

This article contains supporting information.

**Table S1**. List of proteins identified with FDR < 0.01 and quantified. The respective sample for each labelling is as indicated in Figure 1A.

**Table S2**. List of proteins differentially regulated in hypoxia, IOX2, and/or VH032 comparing to DMSO. Proteins were selected at FDR < 0.01 and relative abundance to control DMSO > 1.3 for each replicate. Uniprot ID and gene name are listed with relative abundance to DMSO control. Legends for each sheet are:

A – List of proteins commonly upregulated in hypoxia, IOX2, and VH032 (as in Table 1)

B – List of proteins upregulated only in hypoxia

C – List of proteins upregulated only in IOX2

D – List of proteins upregulated only in VH032

E – List of proteins upregulated in hypoxia and IOX2, but not VH032

F – List of proteins upregulated in hypoxia and VH032, but not IOX2

G – List of proteins upregulated in IOX2 and VH032, but not hypoxia

H – List of proteins commonly downregulated in hypoxia, IOX2, and VH032

I – List of proteins downregulated only in hypoxia

J – List of proteins downregulated only in IOX2

K – List of proteins downregulated only in VH032

L – List of proteins downregulated in hypoxia and VH032, but not IOX2

## Acknowledgements

We would like to thank Wenzhang Chen, Abdel Atrih and Douglas Lamont from the Dundee FingerPrints Proteomic facility for support in sample preparation, processing, and analysis of TMT-labelling mass spectrometry data; Hao Jiang (Dundee) for advice with mass spectrometry data submission; and Alan Fairlamb for discussions.

## Funding and additional information

This work was supported by the Wellcome Trust through a PhD Studentship to JF [102398], by the European Research Council through a starting grant to AC [ERC-2012-StG-311460 DrugE3CRLs], and by Cancer Research UK through a Senior Fellowship to SR [C99667/A12918].

